# Nuclear deformation and anchorage defect induced by DCM mutants in lamin A

**DOI:** 10.1101/611665

**Authors:** Manindra Bera, Rinku Kumar, Bidisha Sinha, Kaushik Sengupta

**Author notes:** These authors contributed equally and the first author was selected on alphabetical order. To whom correspondence may be addressed, Kaushik Sengupta, Biophysics & Structural Genomics Division, Saha Institute of Nuclear Physics, 1/AF, Bidhannagar, Kolkata-700064, India, (Ext. 3504).

## Abstract

Dilated Cardiomyopathy (DCM) is one of the different types of laminopathies caused by the mutations in A-type lamins in somatic cells. The involuntary cyclic stretching of cardiac muscle cells, as observed in normal physiological conditions is perturbed in DCM which afflict patients globally. As A-type lamins are principal components in nuclear mechanics, we have investigated the effect of the DCM causing mutants- K97E, E161K and R190W on nuclear stretching and deformation by static and dynamic strain inducing experiments. All the mutants exhibited differential nuclear structural aberrations along with a tilt in the nuclear axis compared to the direction of the cell axis and a significant decrease in the lamina thickness which reflected the lower mechanical rigidity. These phenotypes could potentially lead to defects in nuclear anchorage to the actin filaments thereby resulting in the misshapen and misaligned nucleus.

## INTRODUCTION

Nuclear lamins which are type V intermediate filament proteins were first visualized to form ~50 nm meshwork underlying the inner nuclear membrane (1). It was considered that lamin A forms 10-nm filaments inside the cell (2–5) until recently when this notion has been revised by the cryo-ET structure of vimentin null mouse embryonic fibroblast suggesting that both A- and B-type lamins form tetrameric 3.5 nm thick filaments inside the nucleus (6). Earlier, it was also reported by the structured illumination microscopy that both lamins form distinct meshwork in the nucleoplasm (7). Lamin A is a principal regulator of nuclear mechanics and the relative abundance of the lamin A is dependent on the tissue types and matrix elasticity (8,9). More than 450 mutations have been discovered in the *LMNA* gene (http://www.umd.be/LMNA/) which produces almost 16 different types of diseases like Dilated Cardiomyopathy (DCM), Emery-Dreifuss Muscular Dystrophy (EDMD) etc. that are collectively coined as laminopathies (10–12). These are tissue-specific in nature predominantly affecting muscle and adipose tissues. The hallmark of laminopathies is the formation of misshapen and even fragile nuclei (13). Two hypotheses are in vogue to explain the role of mutant lamin A protein in the pathogenesis of laminopathies. The gene regulation hypothesis correlated the occurrence of aberrant gene expression due to differential transcriptional regulation by lamin A while the structural hypothesis focuses on the perturbation effect of mutant lamin A on higher order assembly of the nuclear lamina leading to fragile nuclei (14,15). Previously, several reports based on AFM indentation, micropipette aspiration and microrheological experiments showed altered nuclear elasticity for lamin A null MEF or cells containing lamin A mutations (16–18). The rapidly growing area of nucleoskeleton-cytoskeleton interactions has led to numerous studies involving the transduction of external mechanical cues to the nucleus. Earlier studies revealed that cytoskeleton maintains the nucleus in a pre-stressed state by aligned actin filaments forming a perinuclear actin cap (19–21). Recently, it has been shown for endothelial cells that central and apical stress fibers play distinct mechanical roles in maintaining coordination between the cell and nuclear shapes (20).

In this report, we studied the effect of nuclear morphology due to dilated cardiomyopathic mutations (K97E, E161K, and R910W) under different physiological strains. These mutations were selected based on their severities of phenotypes in patients. The phenotype of this disease is characterized by the cardiac arrhythmia with acute conduction defects and myocardial infarction that can lead to sudden death (22). We took two different approaches to observe the effect of the laminopathic mutations (human origin) in the mouse myoblast C2C12. Cells were stretched by applying cyclic strain (dynamic strain) on the PDMS membrane to mimic the physiological state of extension and relaxation of muscle and cardiac cells. Secondly, cells were grown on the different micropatterned surfaces which exert static deformation force (static strain) on the cytoskeleton via cell adhesion molecules (23,24) and ultimately perturbs the lamin A network assembly. In both the cases, we observed differential nuclear deformations, anchorage defect characterized by a tilt in nuclear axis about actin axis and variation in the lamina thickness which suggested a reduction in nuclear rigidity and integrity. These results also suggest a probable explanation behind the formation of elongated and misshapen nuclei in cardiomyopathy caused due to the *LMNA* mutations.

## RESULTS AND DISCUSSION

### Differential impairment of the nucleo-actin axis on the micropatterned substrate

The actin cytoskeleton is known to mechanically couple with the nuclear lamins through **L**inker of **N**ucleoskeleton and **C**ytoskeleton (LINC) complexes (25). We measured the alignment of the nucleus with the cell’s orientation vector in cells expressing WT and mutant Lamins A/C. Usually, the orientation vector of the cell reflects closely the orientation of actin stress fibers (26). However, well-spread cells often had random orientations of stress fibers as is evident from cells on non-patterned glass (Fig. 1 A, left). To be able to control this variation, we used substrates in which the adhesion area was patterned by photomasks (Fig. 1 A, right). Two rectangular patterns denoted by RA and RB with aspect ratios 3, 2 and spread areas 1200, 450 μm^2^ respectively, were chosen. These aspect ratios were chosen based on a similar ratio (1:3) (27)of hPSC-derived monolayers of cardiomyocytes. To quantify the alignment of the nucleus with the cell, the cell and nucleus outlines were fitted with ellipses and the absolute angle between their major axes noted as orientation angle (Fig. 1 B) for cells expressing the GFP tagged versions of lamin A (Fig. 1 C). For perfect alignment of the nucleus with the cell axis, the expected orientation angle is 0°, while for randomly oriented nuclei, the angles are expected to vary from 0° to 90 °, therefore averaging at 45° (dashed line, Fig. 1 D). As suspected, the non-uniform shapes in non-patterned cells resulted in angles (~21°) implying low alignment (Fig. 1 D). There was also no difference in orientation angles in mutant lamins compared to wild type lamin A (denoted by WT) (Fig1 D). However, using micropatterns, we first observed lowered angles (~10°) for WT as expected from the uniformity of the shape (Fig. 1 D). Secondly, we found that all mutants exhibited the alignment of nucleus axis to cell axis within the range 24 ° – 37 ° (Fig. 1 D, Supporting Table1) rendering it closer to the value expected from random orientation (45°, Fig. 1 D, dashed line). However, no particular parameter of nuclear shape (eccentricity, aspect ratio, circularity, – Fig. 1 D) was significantly affected by the mutations. This strongly suggested the lack of nuclear alignment to cell-axis presumably due to a loss of mechanical coupling between the nucleus and cytoplasmic actin network (Fig. 1 D, *top*). Furthermore, we observed the lamin A aggregation inside the nucleoplasm and quantified the size in terms of area. K97E and R190W transfected nuclei showed significantly larger aggregates, ~14 and 12 μm^2^ respectively. WT and E161K nuclei showed smaller aggregates compared, ~2.5 and 8.3 μm^2^ respectively on the RB micropatterned surface. But on the RA pattern, E161K and R190W nuclei produced smaller aggregates compared to RB, ~4.6 and 5.5 μm^2^ respectively (shown in Supplementary Fig1). The size of aggregates for WT and K97E were unchanged, ~2 and 15 μm^2^ respectively. The aggregate may constitute other subtypes of endogenous lamin also because of its homo- and hetero-polymerization nature but we only monitored GFP-fluorescence of the exogenous lamin A. Since all these experiments were performed by transient transfection, that enhance the possibility of overexpression of protein lead to misfolded protein and aggregation. We ruled out this possibility by measuring the non-significant change in the amount of lamin upon transfection through western blotting (shown in Supplementary Fig 3).

**Figure 1.**
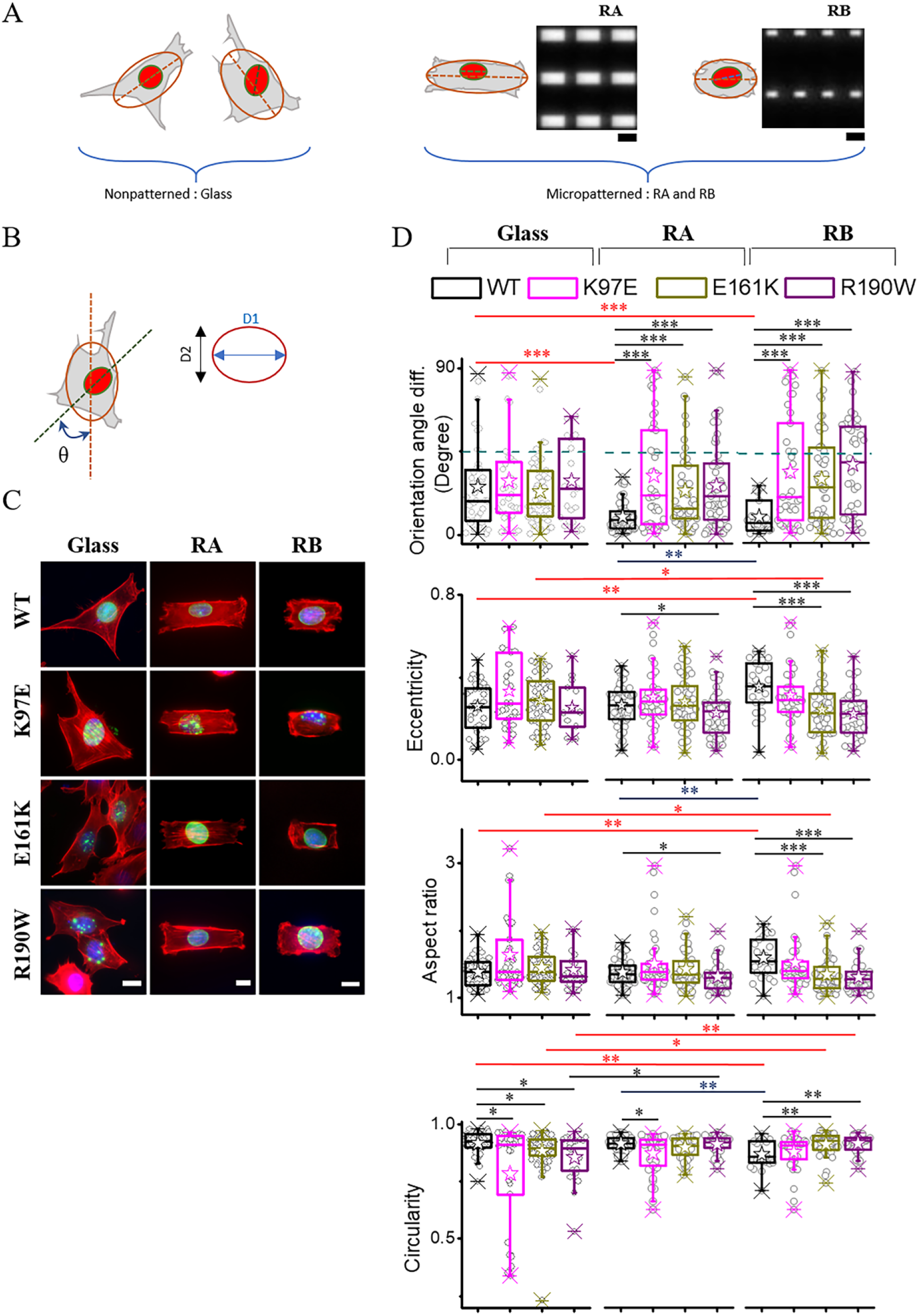
Lamin A mutations lead to the altered alignment of the nucleus with the cell axis. (A) Schematic representations depicting variations possible in the nucleus and cell shape along with nucleus alignment with cell axis in non-patterned (grown on glass) (left) and patterned cells (right). Ellipses are fits to cell-outline and nucleus-outline; the corresponding major axes represented as dashed lines. Wide-field transmission images of patterns on photomasks used to create micropatterns – RA (60 μm × 20 μm) and RB (30 μm × 15 μm) are shown here. Scale bar, 50 μm. (B) Schematic representation of cell to nuclear orientation angle (θ) and major (D1) and minor (D2) diameters of nucleus used for calculating nuclear eccentricity and aspect ratio. (C) Representative images of cells expressing GFP-Lamin A/C (green) mutants stained with DAPI (blue, DNA) and Phalloidin Alexa Fluor 568 (red, F-actin). Images show basal plane for actin and the maximum intensity projection for DAPI and GFP-Lamin A/C. Scale, 10 μm. (D) Orientation angle difference (θ), eccentricity, aspect ratio, and circularity of nucleus measured from three different conditions Glass (*left column*), RA pattern (*middle column*), RB pattern (*right column*). The asterisk marks significance (p < 0.001) difference when compared with WT cells grown on the glass. n = 27, 32, 52, 21 cells for glass from one experiment, n = 70, 41, 48, 55 cells for RA in three independent experiment, n = 41, 37, 52, 36 cells for RB in two independent experiment. * p < 0.05, ** p < 0.01, *** p < 0.001, Wilcoxon based Mann-Whitney U test were performed for statistical testing.

### Nuclear deformations after the cell stretching

In cardiac muscle tissues, the cells continuously experience the cyclic and uniaxial stretching. Hence, the cells also reorient themselves towards the stretching axis (28,29). Zimmermann et al. stretched neonatal rat heart cells up to 10% at 2 Hz frequency to generate engineered heart tissue (30) and Yu et al. showed that 10% static stretching can insert new sarcomere in the neonatal rat (31). The normal resting heart rate in infant and in the case of tachycardia in adult heart rate can be more than 2 Hz (32,33). We intended to test the impact of chronic mechanical perturbation (cyclic stretching) on lamin A-dependent nuclear morphology by mimicking physiological frequency of extension and relaxation of muscles and cardiac cells. We performed the mechanical stretching experiments on the wild-type (WT) and DCM-causing mutant lamin A transfected C2C12 cells. The cells were grown on the PDMS membrane and the membrane was stretched cyclically up to 10% for 2.5 hr. at a frequency of 2 Hz. We measured how the mutants differed from the WT in shape parameters and orientation of nucleus with and without cyclic stretching. In the absence of cyclic stretch, WT and mutant cells grown on PDMS displayed non-significant differences in most (8/12) parameters except for the observations that R190Wwas less aligned than WT while E161K resulted in nuclei with higher eccentricity and aspect ratio and lower circularity. However, after cyclic stretch was imparted most (10/12) parameters were found to be different – especially the ones quantifying nuclei shape changes. Eccentricity and aspect ratio of all three mutants increased on stretching in contrast to E161K which showed an increase even in the absence of stretching. Correspondingly, the circularity of all three mutants decreased on cyclic stretching in contrast to only E161K showing a decrease in the absence of stretching. It must be emphasized that, K97E showed a tendency to deviate from the WT in mechanically unperturbed condition (Fig. 2). On cyclic stretching nuclear deformation of K97Ewas seen to largest among all mutants. Thus, we quantified alignment of the nucleus with the cell body (orientation angle difference) by employing micropatterning to reduce the initial heterogeneity of cell shape in the population (reduced standard deviation of θ, Supporting Table 1). Here, we could elucidate that lamin A mutations may lead to weaker coupling of the nucleus shape to the cell’s shape (Supporting Table 2). Next, we demonstrated that when these mutants undergo mechanical perturbation, they are less resistant to shape changes and therefore underwent deformations, unlike WT nuclei that did not show any significant alteration in nucleus-to-cell alignment or shape on cyclic stretching (Supporting Table 3). We have calculated the meshwork size of lamin A for WT and R190W (Supplementary Fig 2). R190W nuclei dilated lamin A meshwork compared to the WT-nuclei which reflected the lower mechanical rigidity of the nuclei. K97E and E161K nuclei produced lamin A aggregates only.

**Figure 2.**
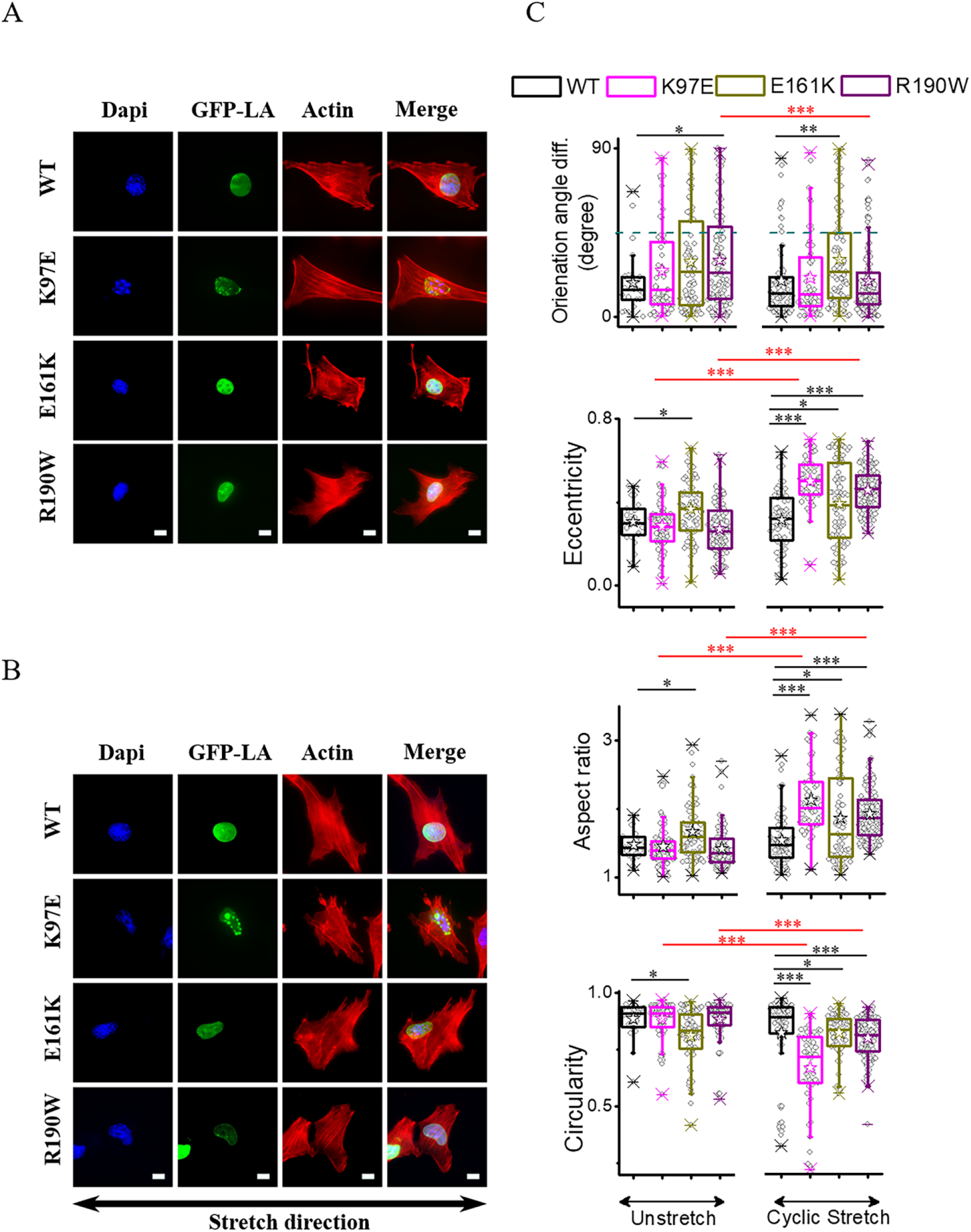
Nuclei of cells with lamin mutations are less resistant to cyclic stretch. Blue, green and red channels in images denote DAPI, GFP-Lamin A transfection and Phalloidin staining in C2C12 cells with **w**ild type (WT) and mutant Lamin A/C (WT, K97E, E161K, R190W). Cells were either (A) grown on PDMS and fixed – without imparting cyclic stretch or (B) grown, imparted cyclic stretch (10% cyclic stretched with 2 Hz frequency for 2.5 h.) and subsequently fixed. Scale, 10 μm. The arrow shows the direction of stretching. (C) Orientation angle difference (θ), eccentricity, aspect ratio, and circularity of nucleus measured from unstretched and stretched conditions. n = 26, 80, 72, 119 cells for unstretched and n = 77, 55, 77, 123 cells for stretched condition in three independent experiment. * p < 0.05 * p < 0.01, ** p < 0.001, Wilcoxon based Mann-Whitney U test were performed for statistical testing.

### Reduction in the lamina thickness due to laminopathic mutation

Nuclear stiffness is primarily determined by nuclear lamina meshwork. We measured lamina thickness which could be the direct readout of stiffness as reported earlier as well (34). We assessed the impact of laminopathic mutations on nuclear lamina thickening behavior. We also included the effect of spreading as reported earlier that lowering in the spread area reduces the stress fiber driven compression of the nucleus and nuclear volume which in turn reduces physical strain on the nucleus (35). We chose two spread area (RA,1200 μm^2^ and RB, 450 μm^2^) which has ~ 3-fold spread area difference. We measured the lamina thickness by structured illumination microscopy (SIM) whose lateral resolution was calculated to be ~120 nm (shown in Supplementary Figure 3). We measured average thickness 0.35 ± 0.03 μm of the WT lamina on the RA patterned that reduces to 0.30 ± 0.03 μm on RB surface (shown in Fig. 3 A-D, left) suggesting thickening of the lamina on highly spread cells. We believed that lamina in the low spread area cells is less stiff thereby resulting in the less aligned network and showed lamina softening behavior. On comparing lamina thickness of WT with R190W and E161K transfected nuclei, we observed that the lamina thickness did not alter for wild-type (~0.3 ± 0.05 μm) and R190W (~0.29± 0.02 μm), however, E161K mutant lamina (~0.23 ± 0.03 μm) showed significant softening behavior compared to WT on the RB surface (Figure 3D). But on RA, the lamina for both E161K and K97E were not clearly visible and both the nuclei showed a significantly large number of aggregates. All the values are tabulated in the supporting table 3. The effect of the geometrical constraint on the K97E-lamin A transfected nuclei were similar. In both cases (RA & RB) for K97E nuclei, the lamina was not clearly visible and nuclei showed larger aggregates. On the spread cell area (RA), the mechanical strain perturbed the lamin A assembly for the mutant nuclei, hence, lamina thickness was significantly decreased or abolished. This lamina softening behavior might explain the lower mechanical rigidity of the nuclei among the cardiomyopathic patients.

**Figure 3.**
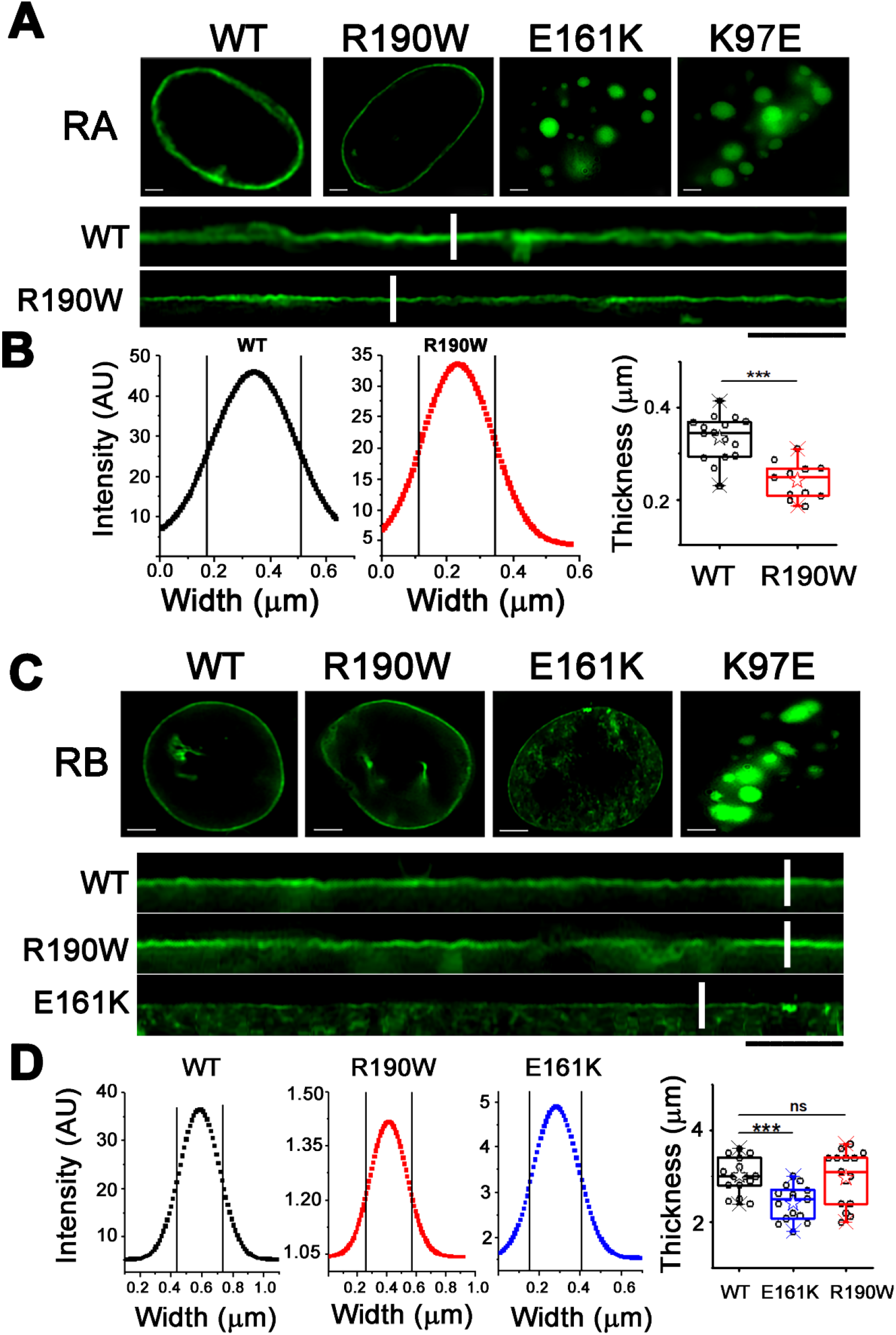
Reduction in the lamina thickness due to laminopathic mutation. eGFP-lamin A (containing wild-type and R190W, E161K & K97E mutations) transfected cells were grown on RA (A) and RB (C) patterned surface. The lower panels of A and C are representing linearization of the nuclear lamina for wild-type and mutant nuclei. The representative FWHMs of the intensity profiles of white the line across the linearized lamina (after Gaussian Fittings) are shown in panel B for RA and in panel D for RB surface. The box plots in the panel B and D represent the FWHM (thickness) of the lamina for RA and RB micropatterned surface respectively. Each dot in the box plot denotes the raw data obtained from each nucleus. The scale bar is 5 μm. ***p<0.001, **p<0.05, ns= not significant.

## CONCLUSION

In this article, we studied the nuclear deformation and misalignments of the nuclear axis compared to the cell axis under different physiological strain. We chose different DCM causing *LMNA* mutants (K97E, E161K, and R190W) and applied external mechanical strain to mimic the physiological condition. In addition, we measured the variation in lamina thickness which plausibly explained the origin of DCM disease and its severity. We observed all the mutations produced significant nuclear deformations and showed significant misalignments of nuclear axes when cyclic stretching was applied. On different micropatterned surfaces, K97E and E161K nuclei produced significantly large lamin A aggregates. Previously, it was reported that no apparent changes in the nuclear lamina occurred for E161K mutation in heart tissue (36). As similarly overexpressed WT nuclei showed less or no visible aggregates, hence the aggregation in the mutant nuclei was due to the overexpression of the improperly assembled mutants lamin A. Previously, it has also been reported that the perinuclear actin and TAN lines anchor the nucleus in pre-stressed condition and help to maintain its orientation (37) and nesprin-1 (an important LINC partner) also helps in tethering the nucleus to actin in C2C12 cells (38). A plausible explanation for impairment of nucleo-actin axis might be the progressive weakening of lamina at points such that the contact nodes with the actin filaments are abrogated thereby resulting in impairment about the axis of the TAN lines. It is important to note that we had previously observed bundling behavior in E161K & R190W which lead to misshapen nuclei with reduced viscoelastic properties. But in the case of the K97E, the lamin A network formation was significantly altered which was reflected on the nuclear elasticity (39,40). Therefore, the cell might not respond to the external mechanical cues effectively which in turn can lead to altered cell response. This might be the reason behind significant nuclear deformations under cyclic stretching. Furthermore, we studied the variation in lamina thickness by SIM. The thickness of the lamina is in the range of ~20-50 nm and electron microscopic study revealed that lamina thickness can vary from ~18 nm to 100 nm in human cartilage due to injury(41). Earlier, Schermmelleh et al. showed by 3D-SIM that thickness of the lamin B1 thread is ~100 nm laterally and ~300 nm axially in C2C12 cells (because of its resolution limit) (42). But in our experiments, lamina thickness was detected around ~0.3 μm which could be due to both overexpression of lamin A and response to mechanical stress. But within the resolution limits (~120 nm) of our experiments, we confirmed the thickening behavior of lamina. We observed that WT lamina thickness slightly increased on the geometrically constrained surface area. But the mutant nuclei showed drastic effect. We could not detect any rim staining of lamin A on the RA surface for the E161K mutant, instead, they formed larger aggregates in the nucleoplasm. The lamina thickness for the R190W mutant nuclei also decreased when cells were grown on the RB surface. We also observed that the total lamin A expression did not change because of the mutation (Supplementary Fig 3). These results suggested that soluble lamin A might get recruited to the lamina in response to an increase in the mechanical strain, but mutant lamin A showed the defect in the lamina assembly. Therefore, R190W nuclei showed less thick lamina and E161K nuclei showed larger lamin A aggregates in the nucleoplasm. Due to mechanical stress, changes in the lamin A structure and the phosphorylation status may lead to a change in the lamina organization (43,44). It is already established that at higher mechanical stress, lamin A accumulates at the nuclear periphery that leads to the increment in nuclear stiffness and at low strain, changes in the phosphorylation status produce more soluble lamin A. Hence, lamin A follows a ‘mechanostat equilibrium’ in the nucleoplasm (45). The change in lamina thickness could be due to the accumulation of the lamin A in the aggregates and the change in the phosphorylation pattern. The increase in WT lamina thickness due to the increase in mechanical stress suggested that changes in the phosphorylation profiles led to the accumulation of the insoluble lamin A to the envelope. But the mutant nuclei lack this ability leading to improper self-assembly and aggregate formation. Ectopic expression of lamin A in the background of the endogenous lamins can significantly contribute to the nuclear stiffness (46). But the background lamin expression was not sufficient to resist the lamina deformation due to different mechanical strain. The aspect ratio of the micropattern was like the hPSC-derived monolayer of the immature cardiomyocytes. The change in the nuclear organization due to the geometrical constraint might be irreversible in the case of cardiomyocytes which might lead to laminopathies. Based on our experimental findings which we have summarized all the results in Supporting Table 1, 2 and 3. We hereby propose that external mechanical cue can alter lamin A meshwork density significantly in presence of laminopathic mutations; lamina shows stress induced thinning behavior and soluble lamin A forms large insoluble aggregates inside the nucleoplasm. These factors cumulatively affect the nuclear rigidity which alters the nuclear shape and mechanics. A plausible model for occurring of laminopathy diseases due to single-point mutation is shown in **Fig 4**. In all cases, we noticed differential alterations in the nuclear shape and the lamin A meshwork for the DCM mutations. The mutant nuclei produced a severe deformation when exposed to the external mechanical strain. In the physiological state also, we can predict that mutation of lamin A in the tissues which are continuously exposed to the external strain can produce significant damage to the nuclear shape and lamin meshwork. Eventually, that can lead to differential gene expression programme and thereby produce the disease.

**Figure 4.**
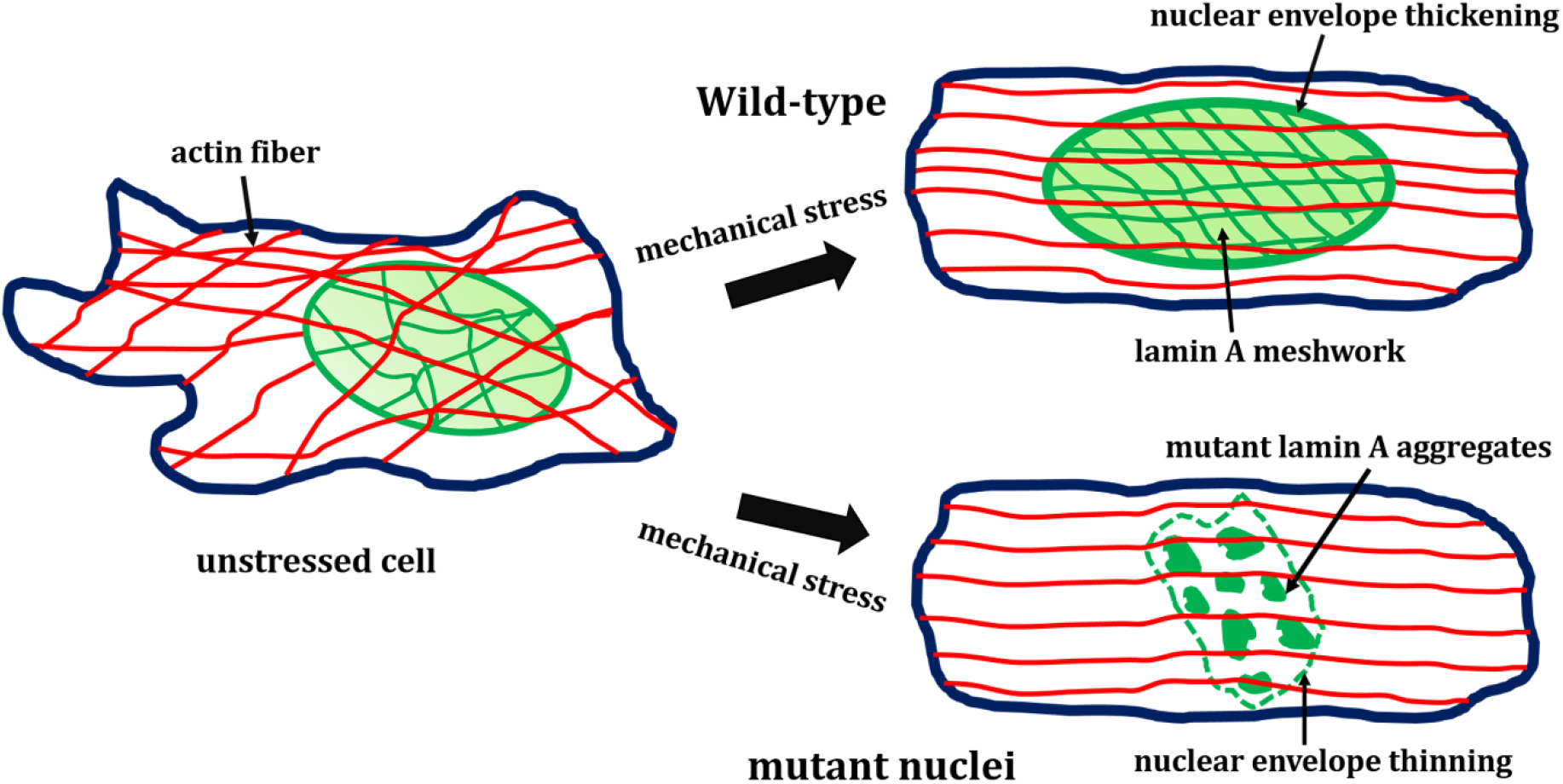
Model for lamin meshwork in the presence of mechanical cue in laminopathic cells. In presence of the external mechanical cue, wild-type lamin A form dense meshwork inside the nucleoplasm and width of the lamina also increases. In the case of laminopathic nuclei, lamin A meshwork density decreases and the lamina also shows thinning behavior in the presence of the same external mechanical force. The nuclear axis also shows large impairment along the acting stress fiber compared to the wild-type nucleus.

## METHODS

### Site-directed mutagenesis

All the mutations were generated using site-directed mutagenesis methods in EGFP-LA plasmid and the details of the primer are reported in Bhattacharjee et al. (40). The mutations were confirmed by the Sanger sequencing method(47).

### Micropatterning glass coverslip

22 × 22 mm glass coverslips were etched in ethanol: acetic acid (19:1) mixture for 30 min, following by ethanol washing and air drying. Dried coverslips were treated with UV Ozone cleaner (Jelight Company, USA) for 5 min and incubated with 0.2 mg/ml PLL-g-PEG (SuSos, Switzerland) solution (prepared in 10 mM HEPES buffer, pH 8.3) for 1 h. Photo-masks (JD, Photo Data, UK) is cleaned with acetone and isopropanol followed by 5 min UV Ozone cleaning. PEG-coated coverslips are attached on the chrome side of already cleaned photo-masks with the help of a drop of water. Excess water was soaked with tissue paper for firm adherence of coverslip to mask. Coverslips attached with photo-mask were illuminated with deep UV for 5 min by placing non-chrome side facing UV lamp. Patterned coverslips were detached by floating into water followed by 45 min incubation in 20 μl fibronectin solution (Sigma) (25 μg/ml solution in NaHCO_3_, pH 8.6). Finally, patterned coverslips were used for cells seeding.

### Transfection and cells seeding

70 % of confluence C2C12 cells were transfected with 3 μg GFP lamin mutants by lipofection (Lipofectamine 3000, Life Technologies) following manufacturer instructions. Post 16 h of transfection, cells were detached from culture dish using 0.02 % EDTA solution (Calbiochem, USA) prepared in cell culture grade PBS (Sigma). 2 × 10^5^ cells were seeded on each patterned coverslips and unattached cells washed off after 30 using warm media. For cell stretching, 10^5^ cells per 500 μl culture media on each PDMS sheet were seeded. After transfection, Lamin A expression was checked mouse monoclonal anti-lamin A/C antibody at 1:1000 dilution.

### Cell Stretching

Custom made cells stretching device was employed for cell stretching as used earlier (ref). Briefly, Silicon elastomer (SYLGRAD 184, Dow Corning) and the curing agent was mixed in 10:1 proportion, an air bubble was removed by centrifuging 5 min at 3000 rpm. 3 ml mixture was spread over each 90 mm dish. Placed vertically to get excess mixture flow away. The dish at this configuration was baked at 60° C to get 100 μm thick PDMS sheet. PDMS sheet was treated 0.5 mg/mlSulfo-SANAPAH (Pierce chemicals) under deep UV for 5 min (UVO cleaner, Jelight, Inc., USA), functionalized with fibronectin (25 μg/ml solution in NaHCO_3_, pH 8.6 Sigma). Fibronectin functionalized coverslips were mounted and custom builds stretchers, GFP Lamin transfected cells were seeded on 25 × 25 mm area encircled with silicon grease. Cells were grown for 24 h on PDMS loaded stretcher inside the incubator. For cyclic cell stretching, PDMS sheet containing C2C12 cells grown in a well was attached to two motors (Physik Instrumente (PI), GmbH and Co KG) using custom-designed adapters. Cyclic stretching at 2 Hz frequency and 10% amplitude was performed with MATLAB for 2.5 h. Stretching was performed inside the cell-culture incubator.

### Cell fixation and microscopy

C2C12 cells either grown on glass or PDMS sheet were fixed using 4% PFA on their respective surfaces for 12 min, following PBS wash. Cells were permeabilization by 0.2 % Triton X-100 solution (Sigma) in PBS, stained with DAPI (2.8 μM, Sigma) and Phalloidin Alexa Fluor 568 (0.35 μM, Molecular probe, Life Technologies) for 2 h at room temperature. Finally, coverslips were mounted on with slides (Sigma) for imaging. In case of cells grown on a PDMS sheet, a number one glass coverslips were first attached on top of the PDS sheet with mounting media (Mowiol, Sigma). The entire stretching device was flipped upside down and captured images using oil objective through the glass. Z-stack (0.5 μm step size) images were captured by Olympus epi-fluorescence microscope (Olympus Corporation Japan) with 100X, 1.49NA objective and sCMOS camera (Orca-Flash 4.0, Hamamatsu Photonics Japan) with pixel size 65 nm.

### Nucleus to cell orientation angle difference and nucleus shape parameter extraction

Image analysis was done using software ImageJ/Fiji ((https://imagej.net/Fiji)). Nucleus (GFP lamin) to cell (actin) orientation angle difference was measured in two-parts, first drawing ROI manually on maximum intensity projection image to outline object, second fitting ROI to an ellipse for measuring orientation angle. Major (D1) and minor (D1) axis of the fitted ellipse on the nucleus was used for eccentricity 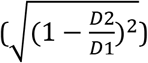 and aspect ratio (D1/D2) measurement. Perimeter and area of object (nucleus) outline on maximum intensity projection image were used for circularity 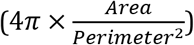 measurement.

### Mesh size and width of the peripheral lamin measurements

Cells were washed three times with ice-cold 1X PBS (pH 7.4) and fixed with 4% paraformaldehyde. Actin was stained with Alexa-561 conjugated phalloidin (Thermo Fisher Scientific) with 1:1000 dilutions. After proper washing with PBS, the coverslips were stained with Vectashield that contained DAPI (Vector Laboratories). The coverslips were sealed with watercolor nail polish. The slides were visualized with NIKON Inverted Research Microscope ECLIPSE TiE with Plan Apo VC 100X oil DIC N2 objective/1.40 NA/1.515 RI with a digital 4X zoom. Images were analyzed using Ni Elements AR Ver 4.13 and Image J (Fiji). The X-Y resolution of N-SIM was calibrated to 120-150 nm using 20 nm FluoSphere beads (Thermo Scientific, F887) in 580 nm laser. We measured the mesh sizes using the area selection tool of Ni-Elements software, but the lines were drawn considering the FWHM method where lower values were considered as the limit of the sides. To measure the lamina thickness, first nuclear rim staining of lamin A were straightened using Image J and then lines were drawn across the linearized lamina and intensity profiles were plotted. From the intensity profile, FHWM was considered to be the thickness or width of the lamina. The thickness of each nucleus was calculated by averaging 10 randomly chosen line profiles and more than 15 separate nuclei were considered for each condition. Sizes of aggregates were measured using an area selection tool of Ni-Elements. The statistics were generated using Origin Pro 8.5 software.

## ACKNOWLEDGMENTS

Authors thank Arikta Biswas for her generous help in the instrument set up.

## AUTHORS CONTRIBUTION

KSG, BS, MB & RK designed all the experiments. RK & MB performed all the experiments. KSG, MB, RK & BS analyzed the data and wrote the paper. KSG conceived the entire project and responsible for finances related to the project.

## REFERENCES

1. Fawcett, D. W. (1966) On the occurrence of a fibrous lamina on the inner aspect of the nuclear envelope in certain cells of vertebrates. The American journal of anatomy 119, 129–145

2. Heitlinger, E., Peter, M., Haner, M., Lustig, A., Aebi, U., and Nigg, E. A. (1991) Expression of chicken lamin B2 in Escherichia coli: characterization of its structure, assembly, and molecular interactions. The Journal of cell biology 113, 485–495

3. Strelkov, S. V., Kreplak, L., Herrmann, H., and Aebi, U. (2004) Intermediate filament protein structure determination. Methods in cell biology 78, 25–43

4. Stuurman, N., Heins, S., and Aebi, U. (1998) Nuclear lamins: their structure, assembly, and interactions. Journal of structural biology 122, 42–66

5. Dechat, T., Pfleghaar, K., Sengupta, K., Shimi, T., Shumaker, D. K., Solimando, L., and Goldman, R. D. (2008) Nuclear lamins: major factors in the structural organization and function of the nucleus and chromatin. Genes & development 22, 832–853

6. Turgay, Y., Eibauer, M., Goldman, A. E., Shimi, T., Khayat, M., Ben-Harush, K., Dubrovsky-Gaupp, A., Sapra, K. T., Goldman, R. D., and Medalia, O. (2017) The molecular architecture of lamins in somatic cells. Nature 543, 261–264

7. Shimi, T., Kittisopikul, M., Tran, J., Goldman, A. E., Adam, S. A., Zheng, Y., Jaqaman, K., and Goldman, R. D. (2015) Structural organization of nuclear lamins A, C, B1, and B2 revealed by superresolution microscopy. Molecular biology of the cell 26, 4075–4086

8. Swift, J., Ivanovska, I. L., Buxboim, A., Harada, T., Dingal, P. C., Pinter, J., Pajerowski, J. D., Spinler, K. R., Shin, J. W., Tewari, M., Rehfeldt, F., Speicher, D. W., and Discher, D. E. (2013) Nuclear lamin-A scales with tissue stiffness and enhances matrix-directed differentiation. Science 341, 1240104

9. Harada, T., Swift, J., Irianto, J., Shin, J. W., Spinler, K. R., Athirasala, A., Diegmiller, R., Dingal, P. C., Ivanovska, I. L., and Discher, D. E. (2014) Nuclear lamin stiffness is a barrier to 3D migration, but softness can limit survival. The Journal of cell biology 204, 669–682

10. Capell, B. C., and Collins, F. S. (2006) Human laminopathies: nuclei gone genetically awry. Nature reviews. Genetics 7, 940–952

11. Bonne, G., Mercuri, E., Muchir, A., Urtizberea, A., Becane, H. M., Recan, D., Merlini, L., Wehnert, M., Boor, R., Reuner, U., Vorgerd, M., Wicklein, E. M., Eymard, B., Duboc, D., Penisson-Besnier, I., Cuisset, J. M., Ferrer, X., Desguerre, I., Lacombe, D., Bushby, K., Pollitt, C., Toniolo, D., Fardeau, M., Schwartz, K., and Muntoni, F. (2000) Clinical and molecular genetic spectrum of autosomal dominant Emery-Dreifuss muscular dystrophy due to mutations of the lamin A/C gene. Annals of neurology 48, 170–180

12. Raffaele Di Barletta, M., Ricci, E., Galluzzi, G., Tonali, P., Mora, M., Morandi, L., Romorini, A., Voit, T., Orstavik, K. H., Merlini, L., Trevisan, C., Biancalana, V., Housmanowa-Petrusewicz, I., Bione, S., Ricotti, R., Schwartz, K., Bonne, G., and Toniolo, D. (2000) Different mutations in the LMNA gene cause autosomal dominant and autosomal recessive Emery-Dreifuss muscular dystrophy. American journal of human genetics 66, 1407–1412

13. Ho, C. Y., Jaalouk, D. E., and Lammerding, J. (2013) Novel insights into the disease etiology of laminopathies. Rare diseases 1, e27002

14. Bertrand, A. T., Chikhaoui, K., Yaou, R. B., and Bonne, G. (2011) Clinical and genetic heterogeneity in laminopathies. Biochemical Society transactions 39, 1687–1692

15. Worman, H. J., and Bonne, G. (2007) “Laminopathies”: a wide spectrum of human diseases. Experimental cell research 313, 2121–2133

16. Radmacher, M. (2007) Studying the mechanics of cellular processes by atomic force microscopy. Methods in cell biology 83, 347–372

17. Lammerding, J., Dahl, K. N., Discher, D. E., and Kamm, R. D. (2007) Nuclear mechanics and methods. Methods in cell biology 83, 269–294

18. Rowat, A. C., Lammerding, J., Herrmann, H., and Aebi, U. (2008) Towards an integrated understanding of the structure and mechanics of the cell nucleus. BioEssays : news and reviews in molecular, cellular and developmental biology 30, 226–236

19. Khatau, S. B., Hale, C. M., Stewart-Hutchinson, P. J., Patel, M. S., Stewart, C. L., Searson, P. C., Hodzic, D., and Wirtz, D. (2009) A perinuclear actin cap regulates nuclear shape. Proceedings of the National Academy of Sciences of the United States of America 106, 19017–19022

20. Versaevel, M., Grevesse, T., and Gabriele, S. (2012) Spatial coordination between cell and nuclear shape within micropatterned endothelial cells. Nature communications 3, 671

21. Mazumder, A., Roopa, T., Basu, A., Mahadevan, L., and Shivashankar, G. V. (2008) Dynamics of chromatin decondensation reveals the structural integrity of a mechanically prestressed nucleus. Biophysical journal 95, 3028–3035

22. Taylor, M. R., Fain, P. R., Sinagra, G., Robinson, M. L., Robertson, A. D., Carniel, E., Di Lenarda, A., Bohlmeyer, T. J., Ferguson, D. A., Brodsky, G. L., Boucek, M. M., Lascor, J., Moss, A. C., Li, W. L., Stetler, G. L., Muntoni, F., Bristow, M. R., Mestroni, L., and Familial Dilated Cardiomyopathy Registry Research, G. (2003) Natural history of dilated cardiomyopathy due to lamin A/C gene mutations. Journal of the American College of Cardiology 41, 771–780

23. Garrison, M. D., McDevitt, T. C., Luginbuhl, R., Giachelli, C. M., Stayton, P., and Ratner, B. D. (2000) Quantitative interrogation of micropatterned biomolecules by surface force microscopy. Ultramicroscopy 82, 193–202

24. Lehnert, D., Wehrle-Haller, B., David, C., Weiland, U., Ballestrem, C., Imhof, B. A., and Bastmeyer, M. (2004) Cell behaviour on micropatterned substrata: limits of extracellular matrix geometry for spreading and adhesion. Journal of cell science 117, 41–52

25. Wang, N., Tytell, J. D., and Ingber, D. E. (2009) Mechanotransduction at a distance: mechanically coupling the extracellular matrix with the nucleus. Nat Rev Mol Cell Biol 10, 75–82

26. Hatch, E. M., and Hetzer, M. W. (2016) Nuclear envelope rupture is induced by actin-based nucleus confinement. The Journal of cell biology 215, 27–36

27. Lundy, S. D., Zhu, W. Z., Regnier, M., and Laflamme, M. A. (2013) Structural and functional maturation of cardiomyocytes derived from human pluripotent stem cells. Stem cells and development 22, 1991–2002

28. Liau, B., Zhang, D., and Bursac, N. (2012) Functional cardiac tissue engineering. Regenerative medicine 7, 187–206

29. Hirt, M. N., Hansen, A., and Eschenhagen, T. (2014) Cardiac tissue engineering: state of the art. Circulation research 114, 354–367

30. Zimmermann, W. H., Melnychenko, I., Wasmeier, G., Didie, M., Naito, H., Nixdorff, U., Hess, A., Budinsky, L., Brune, K., Michaelis, B., Dhein, S., Schwoerer, A., Ehmke, H., and Eschenhagen, T. (2006) Engineered heart tissue grafts improve systolic and diastolic function in infarcted rat hearts. Nature medicine 12, 452–458

31. Yu, J. G., and Russell, B. (2005) Cardiomyocyte remodeling and sarcomere addition after uniaxial static strain in vitro. The journal of histochemistry and cytochemistry : official journal of the Histochemistry Society 53, 839–844

32. Fleming, S., Thompson, M., Stevens, R., Heneghan, C., Pluddemann, A., Maconochie, I., Tarassenko, L., and Mant, D. (2011) Normal ranges of heart rate and respiratory rate in children from birth to 18 years of age: a systematic review of observational studies. Lancet 377, 1011–1018

33. Pinto, D. S., Ho, K. K., Zimetbaum, P. J., Pedan, A., and Goldberger, A. L. (2003) Sinus versus nonsinus tachycardia in the emergency department: importance of age and heart rate. BMC cardiovascular disorders 3, 7

34. Buxboim, A., Irianto, J., Swift, J., Athirasala, A., Shin, J. W., Rehfeldt, F., and Discher, D. E. (2017) Coordinated increase of nuclear tension and lamin-A with matrix stiffness outcompetes lamin-B receptor that favors soft tissue phenotypes. Molecular biology of the cell 28, 3333–3348

35. Kim, D. H., Li, B., Si, F., Phillip, J. M., Wirtz, D., and Sun, S. X. (2015) Volume regulation and shape bifurcation in the cell nucleus. J Cell Sci 128, 3375–3385

36. Mewborn, S. K., Puckelwartz, M. J., Abuisneineh, F., Fahrenbach, J. P., Zhang, Y., MacLeod, H., Dellefave, L., Pytel, P., Selig, S., Labno, C. M., Reddy, K., Singh, H., and McNally, E. (2010) Altered chromosomal positioning, compaction, and gene expression with a lamin A/C gene mutation. PloS one 5, e14342

37. Maninova, M., Iwanicki, M. P., and Vomastek, T. (2014) Emerging role for nuclear rotation and orientation in cell migration. Cell adhesion & migration 8, 42–48

38. Espigat-Georger, A., Dyachuk, V., Chemin, C., Emorine, L., and Merdes, A. (2016) Nuclear alignment in myotubes requires centrosome proteins recruited by nesprin-1. Journal of cell science 129, 4227–4237

39. Banerjee, A., Rathee, V., Krishnaswamy, R., Bhattacharjee, P., Ray, P., Sood, A. K., and Sengupta, K. (2013) Viscoelastic behavior of human lamin A proteins in the context of dilated cardiomyopathy. PloS one 8, e83410

40. Bhattacharjee, P., Banerjee, A., Banerjee, A., Dasgupta, D., and Sengupta, K. (2013) Structural alterations of Lamin A protein in dilated cardiomyopathy. Biochemistry 52, 4229–4241

41. Ghadially, F. N., Dick, C. E., and Lalonde, J. M. (1980) Thickening of the nuclear fibrous lamina in injured human semilunar cartilages. Journal of anatomy 131, 717–722

42. Schermelleh, L., Carlton, P. M., Haase, S., Shao, L., Winoto, L., Kner, P., Burke, B., Cardoso, M. C., Agard, D. A., Gustafsson, M. G., Leonhardt, H., and Sedat, J. W. (2008) Subdiffraction multicolor imaging of the nuclear periphery with 3D structured illumination microscopy. Science 320, 1332–1336

43. Bera, M., Kotamarthi, H. C., Dutta, S., Ray, A., Ghosh, S., Bhattacharyya, D., Ainavarapu, S. R., and Sengupta, K. (2014) Characterization of unfolding mechanism of human lamin A Ig fold by single-molecule force spectroscopy-implications in EDMD. Biochemistry 53, 7247–7258

44. Buxboim, A., Swift, J., Irianto, J., Spinler, K. R., Dingal, P. C., Athirasala, A., Kao, Y. R., Cho, S., Harada, T., Shin, J. W., and Discher, D. E. (2014) Matrix elasticity regulates lamin-A,C phosphorylation and turnover with feedback to actomyosin. Current biology : CB 24, 1909–1917

45. Osmanagic-Myers, S., Dechat, T., and Foisner, R. (2015) Lamins at the crossroads of mechanosignaling. Genes & development 29, 225–237

46. Zwerger, M., Jaalouk, D. E., Lombardi, M. L., Isermann, P., Mauermann, M., Dialynas, G., Herrmann, H., Wallrath, L. L., and Lammerding, J. (2013) Myopathic lamin mutations impair nuclear stability in cells and tissue and disrupt nucleo-cytoskeletal coupling. Human molecular genetics 22, 2335–2349

47. Sanger, F., and Coulson, A. R. (1975) A rapid method for determining sequences in DNA by primed synthesis with DNA polymerase. Journal of molecular biology 94, 441–448

