## Supplementary Information for "Nuclear deformation and anchorage defect induced by DCM mutants in lamin A"

3 Present address: Department of Cell Biology, Yale School of Medicine, New Haven, CT, USA 06510

**Running title:** Nuclear deformations in cardiomyopathic nuclei

**Keywords:** Deformation; Stress; Cardiomyopathic; Nuclei

*To whom correspondence may be addressed

Kaushik Sengupta

Biophysics & Structural Genomics Division, Saha Institute of Nuclear Physics, 1/AF, Bidhannagar, Kolkata-700064, India,

Phone- 033 2337 5345 (Ext. 3504)

### Supplementary Fig 1.

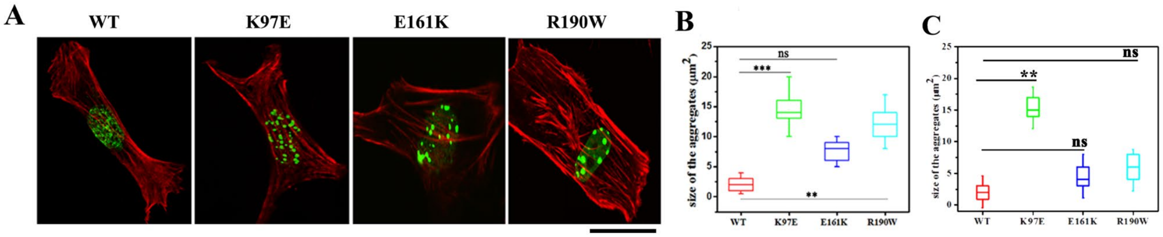

**Lamin A aggregation formation on the micropatterned surface** (A) Different lamin A transfected C2C12 cells were cultured on the rectangular micropatterned surface (RB). Actin was stained with Phalloidin (red) and lamin A/C proteins were in the eGFP-C1 construct (green). The scale bar is 15 μm. The sizes of the aggregates on RB and RA micropatterned surface are shown in B and C respectively. ***p<0.001, **p<0.05, ns= not significant.

**Supplementary Fig 2.**

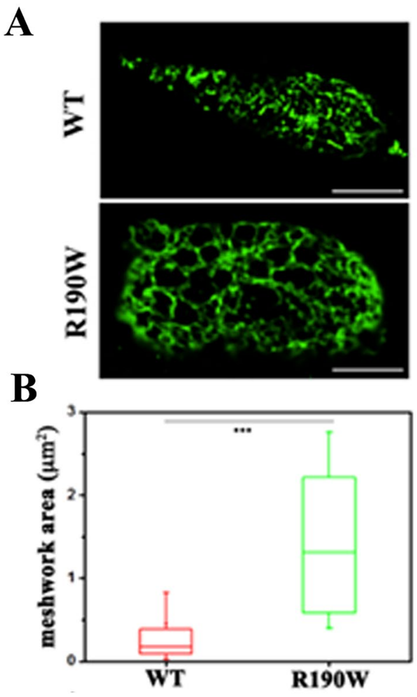

**Lamin A meshwork dilation in response to cyclic stretching.** Representative images of lamin A meshwork after 10% cyclic stretching at 2 Hz frequency. The scale bar is 5 μm. The meshwork box chart in shown in B. Meshwork size of the R190W mutants was larger compared to the wild-type lamin A meshwork***p<0.001, n= 20 nuclei.

**Supplementary Figure 3**

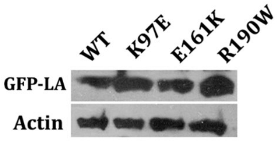

**Expression of the lamin A.** lamin A was probed with anti-lamin A/C mouse monoclonal antibody.

### Supplementary Fig 4.

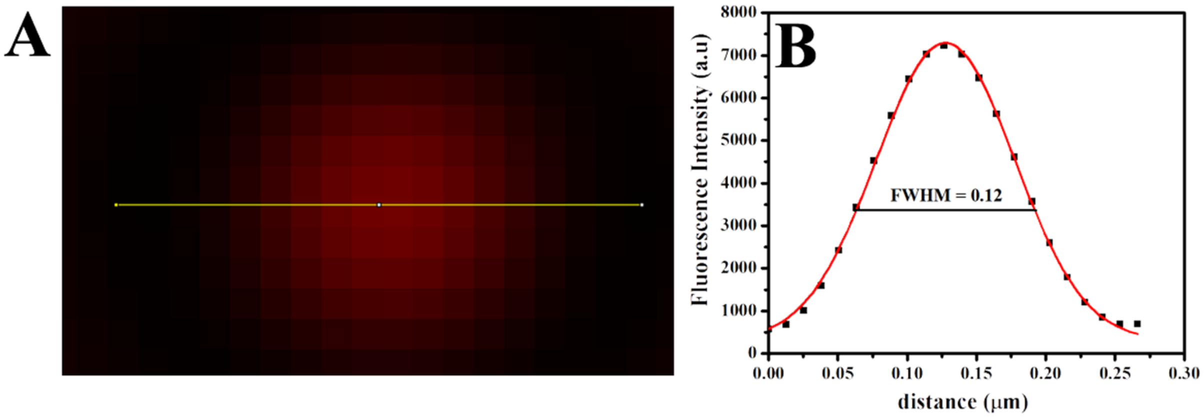

**Supplementary Fig4. Intensity profile of 20 nm fluorescent beads**. A) 20 nm FluoSphere (580/605) in N-SIM. B) the intensity profile along the line (drawn in A). Black dots represent the intensity whereas the red solid line denotes the Gaussian fitting. The calculated FWHM is 0.12 μm.

**Supporting table 1**

| Figure1D | | | | | | | | |
|---|---|---|---|---|---|---|---|---|
| **Parameters Glass** | **Conditions** | **Day** | **Sample size (Cells)** | **Mean** | **Median** | **SD** | **SEM** | **p-value (w.r.t. WT)** |
| Orientation angle (degree) | WT | Day 1 | 39 | 25.87 | 18.33 | 22.53 | 3.61 | |
| | K97E | | 34 | 29.21 | 23.10 | 24.98 | 4.28 | NS |
| | E161K | | 53 | 23.24 | 16.68 | 18.14 | 2.49 | NS |
| | R190W | | 21 | 28.97 | 24.84 | 22.11 | 4.82 | NS |
| Eccentricity | WT | Day 1 | 39 | 0.26 | 0.27 | 0.12 | 0.02 | |
| | K97E | | 34 | 0.33 | 0.27 | 0.17 | 0.03 | NS |
| | E161K | | 53 | 0.29 | 0.29 | 0.12 | 0.02 | NS |
| | R190W | | 21 | 0.26 | 0.24 | 0.13 | 0.03 | NS |
| Aspect ratio | WT | Day 1 | 39 | 1.38 | 1.379 | 0.22 | 0.04 | |
| | K97E | | 34 | 1.64 | 1.410 | 0.55 | 0.09 | NS |
| | E161K | | 53 | 1.44 | 1.386 | 0.25 | 0.03 | NS |
| | R190W | | 21 | 1.41 | 1.319 | 0.29 | 0.06 | NS |
| Circularity | WT | Day 1 | 39 | 0.92 | 0.927 | 0.05 | 0.01 | |
| | K97E | | 34 | 0.78 | 0.911 | 0.23 | 0.04 | 0.036 |
| | E161K | | 53 | 0.88 | 0.896 | 0.10 | 0.01 | 0.034 |
| | R190W | | 21 | 0.86 | 0.895 | 0.11 | 0.02 | 0.015 |
| **Parameters RA** | **Conditions** | **Day** | **Sample size (Cells)** | **Mean** | **Median** | **SD** | **SEM** | **p-value (w.r.t. WT)** |
| Orientation angle (degree) | WT | Day 1,2,3 | 70 | 9.50 | 7.80 | 7.94 | 0.95 | |
| | K97E | | 41 | 31.66 | 21.05 | 28.28 | 4.42 | 4.3E-05 |
| | E161K | | 48 | 23.58 | 14.91 | 21.24 | 3.07 | 3.4E-05 |
| | R190W | | 55 | 26.40 | 20.64 | 22.69 | 3.06 | 9.1E-06 |

| | | | | | | | | |
|---|---|---|---|---|---|---|---|---|
| Eccentricity | WT | Day 1,2,3 | 70 | 0.26 | 0.27 | 0.10 | 0.01 | |
| | K97E | | 41 | 0.30 | 0.28 | 0.14 | 0.02 | NS |
| | E161K | | 48 | 0.28 | 0.26 | 0.13 | 0.02 | NS |
| | R190W | | 55 | 0.23 | 0.23 | 0.11 | 0.02 | 0.034 |
| Aspect ratio | WT | Day 1,2,3 | 70 | 1.38 | 1.39 | 0.20 | 0.03 | |
| | K97E | | 41 | 1.51 | 1.39 | 0.44 | 0.07 | NS |
| | E161K | | 48 | 1.37 | 1.28 | 0.28 | 0.05 | NS |
| | R190W | | 55 | 1.32 | 1.30 | 0.21 | 0.03 | 0.034 |
| Circularity | WT | Day 1,2,3 | 70 | 0.92 | 0.92 | 0.03 | 0.00 | |
| | K97E | | 41 | 0.88 | 0.92 | 0.08 | 0.01 | 0.026 |
| | E161K | | 48 | 0.90 | 0.92 | 0.05 | 0.01 | 0.079 |
| | R190W | | 55 | 0.92 | 0.93 | 0.04 | 0.01 | NS |
| **Parameters RB** | **Conditions** | **Day** | **Sample size (Cells)** | **Mean** | **Median** | **SD** | **SEM** | **p-value (w.r.t. WT)** |
| Orientation angle (degree) | WT | Day 1,2 | 23 | 9.11 | 5.81 | 9.16 | 1.91 | |
| | K97E | | 37 | 33.55 | 20.03 | 29.49 | 4.78 | 4.8E-04 |
| | E161K | | 52 | 29.70 | 26.41 | 24.03 | 3.33 | 1.1E-04 |
| | R190W | | 36 | 37.57 | 39.84 | 25.98 | 4.33 | 1.9E-05 |
| Eccentricity | WT | Day 1,2 | 23 | 0.35 | 0.35 | 0.13 | 0.03 | |
| | K97E | | 37 | 0.31 | 0.29 | 0.14 | 0.02 | NS |
| | E161K | | 52 | 0.24 | 0.22 | 0.12 | 0.02 | 2.7E-04 |
| | R190W | | 36 | 0.22 | 0.22 | 0.11 | 0.02 | 1.9E-04 |
| Aspect ratio | WT | Day 1,2 | 23 | 1.60 | 1.55 | 0.30 | 0.01 | |
| | K97E | | 37 | 1.46 | 1.40 | 0.29 | 0.01 | NS |
| | E161K | | 52 | 1.34 | 1.28 | 0.25 | 0.01 | 3.5E-04 |
| | R190W | | 36 | 1.32 | 1.29 | 0.22 | 0.01 | 3.7E-04 |
| Circularity | WT | Day 1,2 | 39 | 0.87 | 0.86 | 0.06 | 0.00 | |

| | | | | | | | | |
|---|---|---|---|---|---|---|---|---|
| | K97E | | 34 | 0.88 | 0.91 | 0.08 | 0.01 | NS |
| | E161K | | 53 | 0.92 | 0.93 | 0.05 | 0.01 | 1.8E-03 |
| | R190W | | 21 | 0.92 | 0.93 | 0.04 | 0.01 | 4.1E-03 |
| **Figure2D** | | | | | | | | |
| **Parameters** | **Conditions** | **Day** | **Sample size (Cells)** | **Mean** | **Median** | **SD** | **SEM** | **p-value (w.r.t. WT)** |
| Orientation angle (degree) | WT(Unstr) | Day 1,2,3 | 26 | 16.35 | 14.71 | 12.06 | 2.36 | |
| | K97E(Unstr) | | 80 | 24.71 | 14.53 | 23.20 | 2.99 | NS |
| | E161K(Unstr) | | 72 | 29.59 | 24.00 | 26.19 | 2.95 | NS |
| | R190W(Unstr) | | 119 | 30.32 | 23.61 | 25.17 | 2.28 | 0.02 |
| | WT(Str) | | 77 | 19.15 | 12.81 | 19.26 | 2.19 | |
| | K97E(Str) | | 55 | 20.95 | 12.37 | 21.50 | 2.90 | NS |
| | E161K(Str) | | 77 | 30.90 | 24.00 | 24.81 | 2.83 | NS |
| | R190W(Str) | | 123 | 19.16 | 12.60 | 19.12 | 1.73 | NS |
| Eccentricity | WT(Unstr) | Day 1,2,3 | 26 | 0.31 | 0.30 | 0.11 | 0.02 | |
| | K97E(Unstr) | | 80 | 0.29 | 0.29 | 0.12 | 0.01 | NS |
| | E161K(Unstr) | | 72 | 0.37 | 0.37 | 0.14 | 0.02 | 0.04 |
| | R190W(Unstr) | | 119 | 0.27 | 0.26 | 0.12 | 0.01 | NS |
| | WT(Str) | | 77 | 0.32 | 0.32 | 0.14 | 0.02 | |
| | K97E(Str) | | 55 | 0.50 | 0.50 | 0.12 | 0.02 | 6.3E-12 |
| | E161K(Str) | | 77 | 0.40 | 0.39 | 0.19 | 0.02 | 0.01 |
| | R190W(Str) | | 123 | 0.46 | 0.46 | 0.10 | 0.01 | 2.8E-12 |
| Aspect ratio | WT(Unstr) | Day 1,2,3 | 26 | 1.48 | 1.43 | 0.23 | 0.05 | |
| | K97E(Unstr) | | 80 | 1.45 | 1.40 | 0.28 | 0.03 | NS |
| | E161K(Unstr) | | 72 | 1.67 | 1.59 | 0.41 | 0.05 | 0.04 |
| | R190W(Unstr) | | 119 | 1.42 | 1.36 | 0.29 | 0.03 | NS |
| | WT(Str) | | 77 | 1.54 | 1.47 | 0.37 | 0.04 | |
| | K97E(Str) | | 55 | 2.13 | 2.02 | 0.50 | 0.07 | 6.3E-12 |
| | E161K(Str) | | 77 | 1.86 | 1.63 | 0.67 | 0.08 | 0.01 |
| | R190W(Str) | | 123 | 1.93 | 1.87 | 0.40 | 0.04 | 2.8E-12 |
| Circularity | WT(Unstr) | Day 1,2,3 | 26 | 0.89 | 0.91 | 0.08 | 0.02 | |

| | | | | | | | | |
|---|---|---|---|---|---|---|---|---|
| | K97E(Unstr) | | 80 | 0.89 | 0.91 | 0.07 | 0.01 | NS |
| | E161K(Unstr) | | 72 | 0.81 | 0.83 | 0.12 | 0.01 | 0.001 |
| | R190W(Unstr) | | 119 | 0.88 | 0.92 | 0.09 | 0.01 | NS |
| | WT(Str) | | 77 | 0.82 | 0.89 | 0.18 | 0.02 | |
| | K97E(Str) | | 55 | 0.67 | 0.72 | 0.17 | 0.02 | 7.4E-10 |
| | E161K(Str) | | 62 | 0.82 | 0.84 | 0.09 | 0.01 | 0.002 |
| | R190W(Str) | | 123 | 0.80 | 0.81 | 0.09 | 0.01 | 1.2E-6 |

Table 2. Details of statistical parameters. Unstr and Str represent unstretched and stretched condition respectively. Mann-Whitney U test were performed for finding statistical significance.

| Figure 1D | | | | | | | | | | | | |
|---|---|---|---|---|---|---|---|---|---|---|---|---|
| **Orientation angle diff.** | | | | | | | | | | | | |
| | **WT_Glass** | **K97R_Glass** | **E161K_Glass** | **R190W_Glass** | **WT_RA** | **K97E_RA** | **E161K_RA** | **R190W_RA** | **WT_RB** | **K97E_RB** | **E161K_RB** | **R190W_RB** |
| **WT_Glass** | | 0.5580 | 0.9959 | 0.5194 | ***1.25E-09*** | 0.0751 | ***8.75E-04*** | **5.91E-03** | ***1.14E-06*** | 0.1242 | ***3.54E-02*** | 0.5550 |
| **K97R_Glass** | | | 0.5524 | 0.8628 | ***3.42E-09*** | ***3.51E-02*** | ***6.91E-04*** | **2.43E-03** | ***1.85E-06*** | 0.0668 | ***1.09E-02*** | 0.2506 |
| **E161K_Glass** | | | | 0.4463 | ***1.20E-12*** | ***3.85E-02*** | ***2.17E-04*** | **1.13E-03** | ***8.86E-08*** | 0.0797 | ***1.39E-02*** | 0.4629 |
| **R190W_Glass** | | | | | ***1.44E-05*** | 0.0719 | ***6.55E-03*** | 0.0207 | ***2.05E-04*** | 0.1227 | ***3.05E-02*** | 0.2304 |
| **WT_RA** | | | | | | ***4.30E-05*** | ***3.49E-05*** | ***9.04E-06*** | 0.4845 | 1.20E-05 | ***1.61E-06*** | ***2.14E-07*** |
| **K97E_RA** | | | | | | | 0.3335 | 0.6093 | ***8.25E-04*** | 0.7684 | 0.9291 | 0.2814 |
| **E161K_RA** | | | | | | | | 0.6223 | ***9.70E-04*** | 0.1681 | 0.2880 | ***2.36E-02*** |
| **R190W_RA** | | | | | | | | | ***3.84E-04*** | 0.4368 | 0.4949 | 0.0586 |
| **WT_RB** | | | | | | | | | | ***4.83E-04*** | ***1.05E-04*** | ***1.99E-05*** |
| **K97E_RB** | | | | | | | | | | | 0.6444 | 0.4588 |
| **E161K_RB** | | | | | | | | | | | | 0.1602 |
| **Eccentricity** | | | | | | | | | | | | |

| | WT_Glass | K97R_Glass | E161K_Glass | R190W_Glass | WT_RA | K97E_RA | E161K_RA | R190W_RA | WT_RB | K97E_RB | E161K_RB | R190W_RB |
|---|---|---|---|---|---|---|---|---|---|---|---|---|
| **WT_Glass** | | 0.1806 | 0.3295 | 0.8673 | 0.9454 | 0.4332 | 0.6373 | 0.1716 | ***9.3E-03*** | 0.2431 | 0.3253 | 0.1246 |
| **K97R_Glass** | | | 0.4930 | 0.2059 | 0.1855 | 0.8118 | 0.3099 | ***1.8E-02*** | ***0.4774*** | 0.9242 | ***3.2E-02*** | ***1.6E-02*** |
| **E161K_Glass** | | | | 0.3498 | 0.2519 | 0.8985 | 0.6123 | ***9.3E-03*** | ***4.8E-02*** | 0.8011 | ***4.7E-02*** | ***9.6E-03*** |
| **R190W_Glass** | | | | | 0.7550 | 0.3587 | 0.5223 | 0.5244 | ***3.6E-02*** | 0.2169 | 0.7772 | 0.3757 |
| **WT_RA** | | | | | | 0.4468 | 0.6672 | ***3.4E-02*** | ***1.6E-03*** | 0.2294 | 0.1522 | ***3.3E-02*** |
| **K97E_RA** | | | | | | | 0.5370 | ***1.1E-02*** | 0.0510 | 0.5582 | ***4.4E-02*** | ***1.2E-02*** |
| **E161K_RA** | | | | | | | | ***4.4E-02*** | ***2.0E-02*** | 0.2875 | 0.1355 | ***3.6E-02*** |
| **R190W_RA** | | | | | | | | | ***1.3E-04*** | ***2.7E-03*** | 0.6105 | 0.8033 |
| **WT_RB** | | | | | | | | | | 0.0900 | ***6.4E-04*** | ***2.0E-04*** |
| **K97E_RB** | | | | | | | | | | | ***1.9E-02*** | ***3.7E-03*** |
| **Aspect ratio** | | | | | | | | | | | | |
| | **WT_Glass** | **K97R_Glass** | **E161K_Glass** | **R190W_Glass** | **WT_RA** | **K97E_RA** | **E161K_RA** | **R190W_RA** | **WT_RB** | **K97E_RB** | **E161K_RB** | **R190W_RB** |
| **WT_Glass** | | 0.1064 | 0.3100 | 0.9567 | 0.9975 | 0.4078 | 0.5588 | 0.1658 | ***5.9E-03*** | 0.2341 | 0.3400 | 0.1203 |
| **K97R_Glass** | | | 0.3364 | 0.1408 | 0.0844 | 0.5231 | 0.1940 | ***4.5E-03*** | ***0.4445*** | 0.7823 | ***1.0E-02*** | ***4.0E-03*** |
| **E161K_Glass** | | | | 0.4218 | 0.3082 | 0.9939 | 0.6389 | ***8.2E-03*** | ***2.7E-02*** | 0.6490 | ***4.3E-02*** | ***9.0E-03*** |
| **R190W_Glass** | | | | | 0.8876 | 0.3721 | 0.6764 | 0.3765 | ***3.3E-02*** | 0.1873 | 0.5592 | 0.2571 |
| **WT_RA** | | | | | | 0.4468 | 0.6672 | ***3.4E-02*** | ***1.6E-03*** | 0.2294 | 0.1522 | ***3.3E-02*** |
| **K97E_RA** | | | | | | | 0.5370 | ***1.1E-02*** | ***5.1E-02*** | 0.5582 | ***4.4E-02*** | ***1.2E-02*** |
| **E161K_RA** | | | | | | | | ***4.4E-02*** | ***2.0E-02*** | 0.2875 | 0.1355 | ***3.6E-02*** |
| **R190W_RA** | | | | | | | | | ***1.3E-04*** | ***2.7E-03*** | 0.6105 | 0.8033 |
| **WT_RB** | | | | | | | | | | 0.0900 | ***6.4E-04*** | ***2.0E-04*** |
| **K97E_RB** | | | | | | | | | | | ***1.9E-02*** | ***3.7E-03*** |
| **E161K_RB** | | | | | | | | | | | | 0.5351 |

| **Circularity** | | | | | | | | | | | | |
|---|---|---|---|---|---|---|---|---|---|---|---|---|
| | **WT_Glass** | **K97R_Glass** | **E161K_Glass** | **R190W_Glass** | **WT_RA** | **K97E_RA** | **E161K_RA** | **R190W_RA** | **WT_RB** | **K97E_RB** | **E161K_RB** | **R190W_RB** |
| **WT_Glass** | | ***3.6E-02*** | ***3.4E-02*** | ***1.5E-02*** | 0.7068 | ***2.8E-02*** | 0.1491 | 0.5908 | ***8.9E-03*** | ***3.3E-02*** | 0.9744 | 0.9662 |
| **K97R_Glass** | | | 0.6987 | 0.9447 | ***6.4E-02*** | 0.6319 | 0.3995 | 0.1187 | 0.9222 | 0.6411 | 0.0525 | ***4.1E-02*** |
| **E161K_Glass** | | | | 0.2190 | ***6.0E-03*** | 0.7604 | 0.3840 | ***3.6E-02*** | 0.1494 | 0.8924 | ***1.1E-02*** | ***1.4E-02*** |
| **R190W_Glass** | | | | | ***9.6E-03*** | 0.4048 | ***7.6E-02*** | ***1.0E-02*** | 0.8233 | 0.4421 | ***4.6E-03*** | ***6.5E-03*** |
| **WT_RA** | | | | | | ***2.6E-02*** | ***7.7E-02*** | 0.9140 | ***1.7E-03*** | ***2.0E-02*** | 0.7718 | 0.5459 |
| **K97E_RA** | | | | | | | 0.3293 | ***4.8E-02*** | 0.6594 | 0.9681 | ***1.8E-02*** | ***1.3E-02*** |
| **E161K_RA** | | | | | | | | 0.2119 | ***4.8E-02*** | 0.3612 | 0.0750 | 0.0771 |
| **R190W_RA** | | | | | | | | | ***6.6E-03*** | ***3.0E-02*** | 0.4755 | 0.5018 |
| **WT_RB** | | | | | | | | | | 0.4748 | ***1.7E-03*** | ***4.0E-03*** |
| **K97E_RB** | | | | | | | | | | | ***1.8E-02*** | ***1.2E-02*** |
| **E161K_RB** | | | | | | | | | | | | 0.9560 |
| Figure2 C | | | | | | | | | | | | |
| **Orientation angle diff.** | | | | | | | | | | | | |
| | **WT_UnStr** | **K97R_UnStr** | **E161K_UnStr** | **R190W_UnStr** | **WT_Str** | **K97E_Str** | **E161K_Str** | **R190W_Str** | | | | |
| **WT_UnStr** | | 0.3948 | 0.0609 | ***2.2E-02*** | 0.8525 | 0.9718 | ***1.3E-02*** | 0.7933 | | | | |
| **K97R_UnStr** | | | 0.3595 | 0.1341 | 0.2548 | 0.3082 | 0.0911 | 0.1844 | | | | |
| **E161K_UnStr** | | | | 0.7179 | ***2.6E-02*** | 0.0726 | 0.5085 | ***1.3E-02*** | | | | |
| **R190W_UnStr** | | | | | ***2.7E-03*** | ***1.3E-02*** | 0.7723 | ***3.5E-04*** | | | | |
| **WT_Str** | | | | | | 0.9632 | ***1.5E-03*** | 0.8925 | | | | |
| **K97E_Str** | | | | | | | ***9.2E-03*** | 0.9608 | | | | |
| **E161K_Str** | | | | | | | | ***3.1E-04*** | | | | |
| **Eccentricity** | | | | | | | | | | | | |

| | WT_UnStr | K97R_UnStr | E161K_UnStr | R190W_UnStr | WT_Str | K97E_Str | E161K_Str | R190W_Str |
|---|---|---|---|---|---|---|---|---|
| **WT_UnStr** | | 0.3842 | ***4.1E-02*** | ***1.0E-01*** | 0.8376 | ***5.8E-09*** | ***5.7E-02*** | ***2.0E-08*** |
| **K97R_UnStr** | | | ***2.7E-04*** | 0.2934 | 0.1676 | ***2.6E-15*** | ***8.9E-04*** | ***1.4E-18*** |
| **E161K_UnStr** | | | | ***3.5E-06*** | ***2.4E-02*** | ***6.9E-08*** | 0.3700 | *5.6E-06* |
| **R190W_UnStr** | | | | | ***2.0E-02*** | ***4.2E-18*** | ***1.5E-05*** | ***5.1E-24*** |
| **WT_Str** | | | | | | ***6.4E-12*** | ***1.3E-02*** | ***2.8E-12*** |
| **K97E_Str** | | | | | | | ***2.1E-03*** | ***7.1E-03*** |
| **E161K_Str** | | | | | | | | ***2.2E-02*** |
| **Aspect ratio** | | | | | | | | |
| | **WT_UnStr** | **K97R_UnStr** | **E161K_UnStr** | **R190W_UnStr** | **WT_Str** | **K97E_Str** | **E161K_Str** | **R190W_Str** |
| **WT_UnStr** | | 0.3842 | ***4.1E-02*** | ***1.0E-01*** | 0.8376 | ***5.8E-09*** | ***5.7E-02*** | ***2.0E-08*** |
| **K97R_UnStr** | | | ***2.7E-04*** | 0.2934 | 0.1676 | ***2.6E-15*** | ***8.9E-04*** | ***1.4E-18*** |
| **E161K_UnStr** | | | | ***3.5E-06*** | ***2.4E-02*** | ***6.9E-08*** | 0.3700 | ***5.6E-06*** |
| **R190W_UnStr** | | | | | ***2.0E-02*** | ***4.2E-18*** | ***1.5E-05*** | ***5.1E-24*** |
| **WT_Str** | | | | | | ***6.4E-12*** | ***1.3E-02*** | ***2.8E-12*** |
| **K97E_Str** | | | | | | | ***2.1E-03*** | ***7.1E-03*** |
| **E161K_Str** | | | | | | | | ***2.2E-02*** |
| **Circularity** | | | | | | | | |
| | **WT_UnStr** | **K97R_UnStr** | **E161K_UnStr** | **R190W_UnStr** | **WT_Str** | **K97E_Str** | **E161K_Str** | **R190W_Str** |
| **WT_UnStr** | | 0.9513 | ***1.0E-03*** | 8.7E-01 | 0.3744 | ***7.5E-10*** | ***2.0E-03*** | ***1.2E-06*** |
| **K97R_UnStr** | | | ***2.4E-06*** | 0.7101 | 0.1502 | ***2.8E-18*** | ***8.9E-06*** | ***4.7E-15*** |
| **E161K_UnStr** | | | | ***3.1E-07*** | ***1.4E-02*** | ***2.2E-07*** | 0.0745 | ***1.1E-01*** |
| **R190W_UnStr** | | | | | ***8.9E-02*** | ***2.9E-19*** | ***8.8E-07*** | ***2.1E-17*** |
| **WT_Str** | | | | | | ***5.0E-09*** | ***4.1E-03*** | ***5.5E-06*** |

| | | | | | | | | |
|---|---|---|---|---|---|---|---|---|
| **K97E_Str** | | | | | | | ***2.5E-01*** | ***1.3E-07*** |
| **E161K_Str** | | | | | | | | ***3.9E-01*** |

Mann-Whitney U test was performed for finding statistical significance in pairs. All p-value less than 0.05 are highlighted in bold and italic font.

**Supporting Table 3: Lamina thickness calculations on micropatterned surface**

| **RA** | | **Mean** | **Median** | **Std dev** | **SEM** | **p-value** |
|---|---|---|---|---|---|---|
| | **WT (n= 17)** | 0.35 | 0.34 | 0.048 | 0.012 | |
| | **R190W (n= 12)** | 0.24 | 0.24 | 0.04 | 0.012 | 2.06E-5 |
| | **E161K (n= 15)** | - | | - | - | - |
| | **K97E (n= 15)** | - | | - | - | - |
| **RB** | **WT (n= 17)** | 0.30 | 0.30 | 0.037 | 0.009 | - |
| | **R190W (n= 17)** | 0.29 | 0.31 | 0.05 | 0.013 | 0.73 |
| | **E161K (n= 17)** | 0.23 | 0.239 | 0.035 | 0.008 | 1.13E-5 |
| | **K97E (n= 17)** | - | | - | - | - |

Lamina thickness measured on the RA and RB surface is reported Figure 3. n denotes the total number of nuclei measured.
